# Automatic identification of intestinal parasites in reptiles using microscopic stool images and convolutional neural networks

**DOI:** 10.1101/2022.02.06.479317

**Authors:** Felipe Grijalva, Bryan Núñez, Alejandra Núñez, Carla Parra, Noel Pérez, Diego Benítez

## Abstract

Captive environments trigger the propagation and multiplication of parasites among different reptile species, thus weakening their immune response and causing infections and diseases. Technological advances of convolutional neural networks have opened a new field for detecting and classifying diseases which have shown great potential to overcome the shortcomings of manual detection performed by experts. Therefore, we propose an approach to identify six captive reptiles parasitic agents (*Ophionyssus natricis, Blastocystis sp, Oxiurdo egg, Rhytidoides similis, Strongyloides, Taenia*) from a microscope stool images dataset. Towards this end, we first use an image segmentation stage to detect the parasite within the image, which combines the Contrast Limited Adaptive Histogram Equalization (CLAHE) technique, the OTSU binarization method, and morphological operations. Then, we carry out a classification stage through Mobilenet CNN under a transfer learning scheme. This method was validated on a stool image dataset containing 3616 images data samples and 26 videos from the six parasites mentioned above. The results obtained indicate that our transfer learning-based approach can learn a helpful representation from the dataset. We obtained an average accuracy of 98.66% across the six parasitic agent classes, which statistically outperformed, at a 95% confidence level, a custom CNN trained from scratch.

## Introduction

Reptile parasitology research has not been fully explored in the scientific literature [1]. Parasites are one of the most common infectious agents and easily spread within wildlife management and care centers [2], which can cause injury or immune suppression in reptiles, increasing the mortality rate or leading to secondary diseases [3]. It has been shown that some types of parasites cause hepatitis in snakes [4]; others affect the behavior and physiology of the lizards, and even some parasites can cause chronic enteritis with edema and hemorrhagic intestinal mucosa [4]. Also, mites in reptiles cause weakness due to blood loss, pneumonia, and even septicemia [5]. The need to control parasitism in captive species is important because they can be a source of transmission of zoonotic diseases, which in turn may be transmitted from animal to human [6]. Manual parasite classification methods involve analyzing stool microscope images by experts. However, this task requires significant effort and time since, in a center where reptiles are sheltered, they may have a very high parasite load of different types, so that it could affect their health. Automatic classification tools for parasitic animal agents can help researchers or keepers to operate much faster and efficiently. Consequently, it is possible to perform faster diagnoses and prepare the adequate deworming protocols to prevent diseases in captive reptiles.

In this context, convolutional neural networks (CNNs) have proven to be a valuable tool to find and classify patterns in images easier than the specialists, which invest long identification times prone to error. CNNs have been studied for several years, achieving good results in image classification tasks, and by using similar models, it is possible to generalize algorithms to solve different types of problems [7]. For example, these machine learning tools have recently been tested in many fields including medicine and biology [8]. There are several factors [9] that influence the effectiveness of CNNs (e.g., architecture, data processing, segmentation). Moreover, the complexity involved in training a CNN from scratch is in general not feasible when the data are insufficient. Therefore, this has led to the use of supporting methods such as data augmentation and transfer learning [10]. The latter uses a previously trained CNN with a dataset and a base task, and then the learned features (network weights) are transferred to a second network to be trained on a new dataset and a target task [11].

In light of this, this paper proposes a new approach for identifying parasitic agents from microscopic stool images that affect reptiles in captivity, through segmentation algorithms, data augmentation strategies, and CNNs under a transfer learning scheme.

### Related work

This section describes several previous studies on similar domains as our approach that employ machine learning techniques. For example, in [12] a model for the detection of adult whitefly (*Bemisia tabaci*) and thrips (*Frankliniella occidentalis*) in greenhouses was proposed. An image acquisition system using adhesive traps allowed the collection of the database. Segmentation was performed using the OTSU algorithm and other digital image processing methods. Finally, the classification was carried out with help of a feed-forward neural network. Similar work is addressed by [13], where several morphological features related to the size and color of the specimens were extracted and analyzed to classify them.

In [14] the authors present a method for the detection and binary classification of cells infected by the malaria parasite. They propose a segmentation stage based on morphological top-hat operators [15], and the classification stage uses different sets of texture and shape features that feed a neural network.

Moreover, in [16], a real-time remote insect trap monitoring system employing IoT and a method for classifying insects based on a Faster region-based CNN (R-CNN) and ResNet 50, applying transfer learning was proposed. The results show that the system could automatically identify insects with 94% accuracy.

Finally, in [17] an approach to classify protozoa and metazoa organisms was proposed. Specifically, they compare discriminant analysis, neural networks and decision trees. They found that the discriminant analysis and the neural network performances were quite similar, while the decision tree technique was less efficient.

Despite these efforts, as far as we know, this is the first attempt to build a machine learning model to specifically classify reptilian parasites using convolutional neural networks from microscopic stool images. For this, as in previous works, we will use traditional digital image processing techniques such as binarization and segmentation [12, 14, 17] in a pre-processing stage to prepare image samples for our classifier. Then, we are going to use a CNN-based classifier under a transfer learning scheme as in [16]. The contributions of this paper are related to (a) the unprecedented study of an end-to-end machine learning model based on image segmentation and CNNs for learning expressive features from microscopic stool images to discriminate reptilian parasites; (b) the comparison of a transfer learning scheme against a custom CNN training from scratch, and (c) the introduction of a new public dataset containing microscopic stool images collected and annotated by experts.

## Materials and methods

Fig 1 shows the block diagram of our proposed approach. The first stage involves collecting images from reptile feces. A veterinarian specialist labeled the images according to the parasite present in them. These images must pass through a segmentation process to reduce background noise to appreciate the parasite as clearly as possible. Also, they must go through a process of data augmentation to deal with the unbalanced classes. The pre-trained MobileNet architecture is used to train a new model on our images in the transfer learning stage. Finally, the trained model predicts the type of parasite for previously unseen reptile feces images.

**Fig 1.**
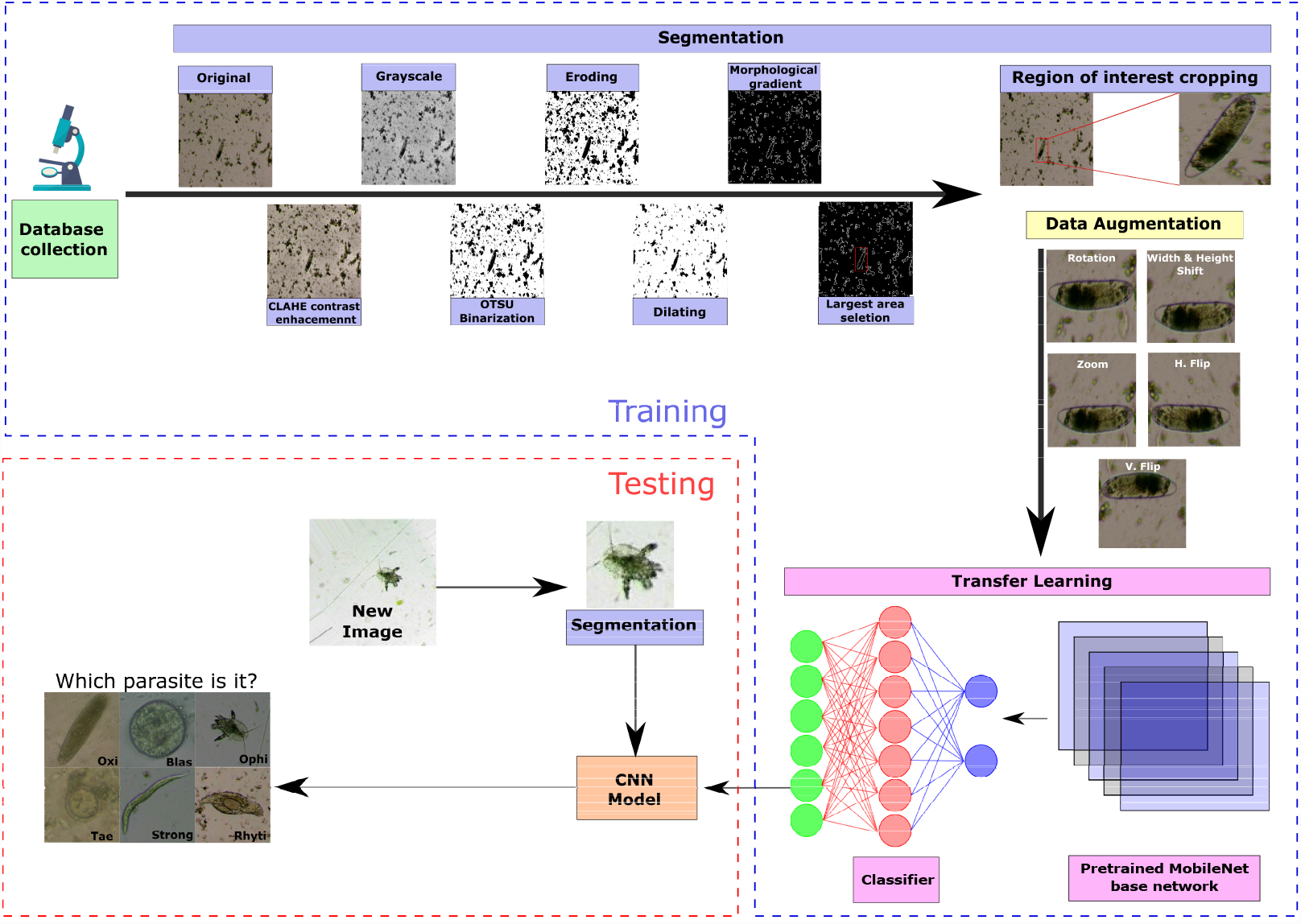
Block diagram of the proposed approach.

### Dataset

The dataset contains images of coproparasitic samples of two orders of reptiles: *Chelonians* (e.g., aquatic, semi-aquatic and terrestrial turtles) and *Squamates* (*Ophidians* such as venomous and non-venomous snakes, and *Saurians* such as lizards, geckos and iguanas) that live in captivity in the Vivarium in Quito (Ecuador)^1^, which houses approximately 350 animals of different species of amphibians and reptiles.

The collection of stool samples was carried out by the indirect method, i.e., from the samples of stool deposited in the areas where each animal is located. The samples were collected with a tongue depressor for each sample to avoid contamination, and then they were stored within vacuum sealed Ziploc bags. These bags were labeled and put into an airtight cooler container. In total, it was collected 118 stool samples.

The stool samples were processed through three stool examinations. Stool tests are the most widely used methods to diagnose parasitic infections. The first was the direct method that investigates fresh stool samples looking for mostly mobile parasitic forms under the microscope. The second one is the flotation technique, which makes the parasitic forms float towards the surface due to their lower density than the solution in which they are immersed. The third is through sedimentation, which uses gravity so that each parasite comes to sediment naturally in a medium with a lower density.

Since some method makes it possible to better identify some parasites than the others from the same sample, each sample was analyzed through a microscope by the three previously described tests. Digital images of parasites were obtained using a digital tactile microscope (Better Scientific Led Q190A-LCD, magnification of X10, X40, and X100) to obtain several images and videos of the parasites. Finally, each image and video were annotated by a veterinarian specialist using the labels shown in Table 1 according to the parasite present in them.

**Table 1.** Parasites database distribution

| Parasites (labels) | Total images | Videos |
| --- | --- | --- |
| Mite - *Ophionyssus natricis* (ophi) | 245 | 2 (1236 frames) |
| *Blastocystis sp* (blas) | 950 | 0 |
| *Oxiurdo egg* (oxi) | 345 | 6 (888 frames) |
| *Rhytidoides similis* (rhyti) | 1072 | 6 (1048 frames) |
| *Strongyloides* (strong) | 648 | 11 (764 frames) |
| *Taenia* (tae) | 356 | 1 (913 frames) |
| **Total** | **3616** | **4849** |

By using the described procedure, a dataset of 3616 images and 26 videos containing 4849 frames from six parasites (see Fig 2 for examples of each parasite) was constructed according to the distribution shown in Table 1. For this study, we extracted all the frames from the 26 videos, which were used only during the training stage of the neural network.

**Fig 2.**
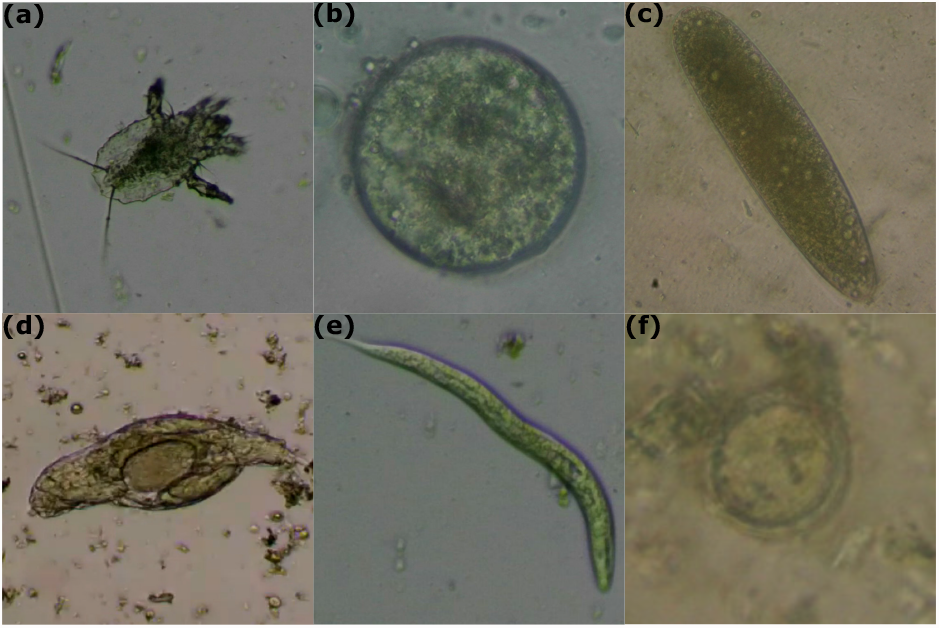
Examples of each parasite. (a) Acaro (*Ophionyssus natricis*); (b) *Blastocystis sp*; (c) *Oxiurdo egg*; (d) *Rhytidoides similis*; (e) *Strongyloides*; (f) *Taenia*.

Since the indirect method employed in this study involves only the collection of samples of stool deposited in the areas where each animal is located and not directly from the animal, ethical approval was not required. The samples were collected by author A. Nuñez specifically for this study.

### Proposed method

#### Region segmentation

Segmentation is a digital image procedure that extracts the region of interest from the original image [18]. The segmentation stage was crucial to achieving good performance in the network training stage since most of the images in the database are very noisy (e.g., as shown in Fig 3a) since the images were taken from animal feces. With this aim, the images were processed by the following steps as depicted in Fig 3.

**Fig 3.**
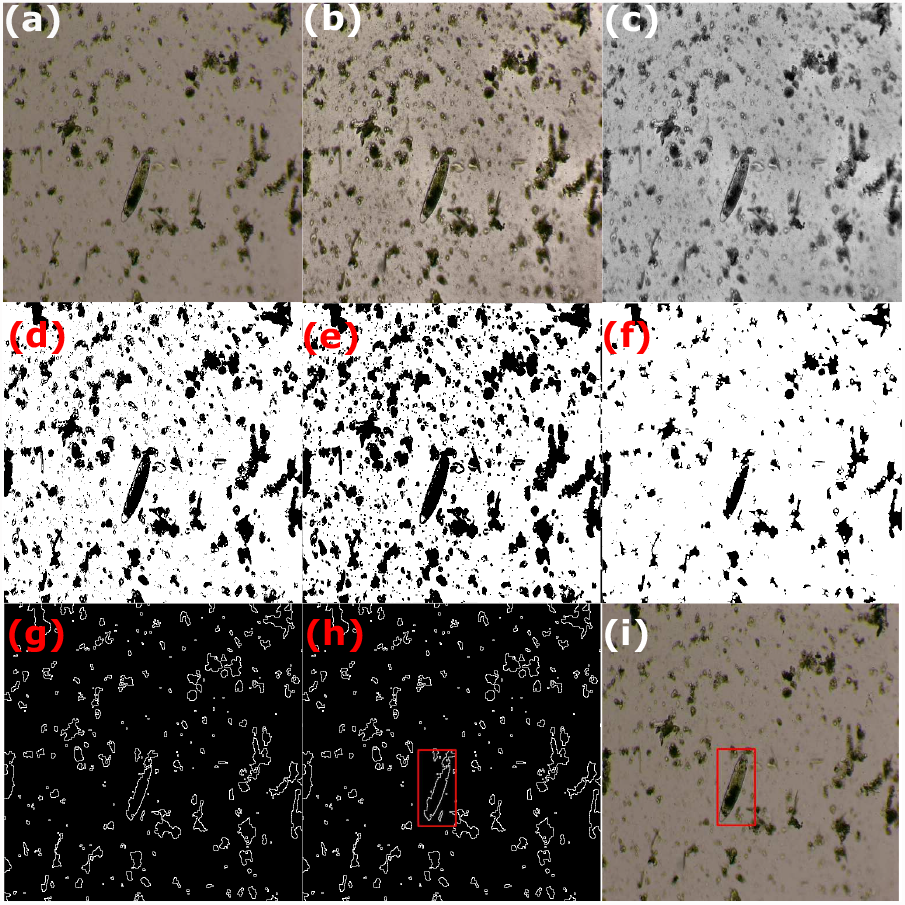
Parasite detection using image processing techniques: (a) original image; (b) image contrast enhancement using CLAHE; (c) color to grayscale image conversion; (d) image binarization using the OTSU method; (e) erode-based morphological operation; (f) dilate-based morphological operation; (g) morphological gradient operation; (h) selection of the largest area (region of interest) in the processed image, and (i) plotting the region of interest in the original image.

- CLAHE (Contrast Limited Adaptive Histogram Equalization) is a digital image processing technique that improves the image contrast without increasing noise. It selects different sections of the image to redistribute their pixel brightness values. As a result, the image contrast is improved while preserving the contours of the objects [19], as shown in Fig 3b.
- After converting the RGB (red, green, blue) color image to grayscale, the image binarization through the Otsu’s method [20] is applied, which determines the most appropriate conversion threshold by minimizing the intra-class variance between two assumed pixel classes (usually, black and white), as shown in Fig 3d.
- Morphological operations [21] (Fig 3e and 3f) are performed by eroding and then dilating the image pixels in order to decrease noise in the image through the kernel

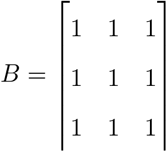

Observe that erosion tends to remove small objects due to debris or garbage so that only substantive objects remain, whereas dilation makes objects such as the parasite more visible.

The morphological gradient (Fig 3g) of an image *I* is then obtained by calculating the difference between the dilation (⊕) and erode (⊖) operations from the previous step using the kernel *B*, according to the following equation

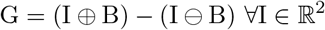

In the resulting image *G*, the contours of the most significant objects are emphasized [22].

- Finally, the areas of possible objects are computed after the morphological gradient operation application. The most significant areas are very likely to contain the target object (parasite). Thus, these areas were cropped and used to feed the neural network classifier, as shown in Fig 3h and 3i.

#### Data augmentation

The objective of data augmentation techniques is to generate new image samples by transforming the original one. Usually, these transformations are affine transformations, i.e., projective transformations that do not move the images’ objects [33]. Thus, they preserve the objects’ collinearity features in the space under analysis. At present, data augmentation is a practical solution to train deep network models which demand a vast amount of image samples, avoiding model overfitting. We used the following augmentation strategies:

- Rotation range: It is the degree range for random rotations. We used a range between -180 to +180 degrees.
- Width shift range: It randomly shifts an image to the left or right by a proportional percentage of the image width. This value was set to 0.2, i.e., 20% of the image width.
- Height shift range: Similar to the previous transformation, but the shift is up or down. This value was set to 0.2.
- Zoom range: It allows for varying the zoom of an image randomly. This value was set to 0.2, i.e., the zoom range lies between 80% to 120%.
- Horizontal flip: It randomly allows an image to be flipped horizontally.
- Vertical flip: It randomly allows an image to be flipped vertically.

Through this process, we augmented the dataset instances so that each class has roughly 1500 images, including original images, augmented images, and video frames to increase the opportunity for a better model’s performance. Since the videos contain parasites in movement, which are visually similar to resulting images from data augmentation procedures, we avoid using data augmentation strategies in the video samples.

Finally, for the rhyti class, the data augmentation algorithms were not used since the instances from this class in conjunction with video frames already reached more than 2000 images. After performing these procedures, 10099 images were obtained with the distribution shown in Table 2.

**Table 2.**
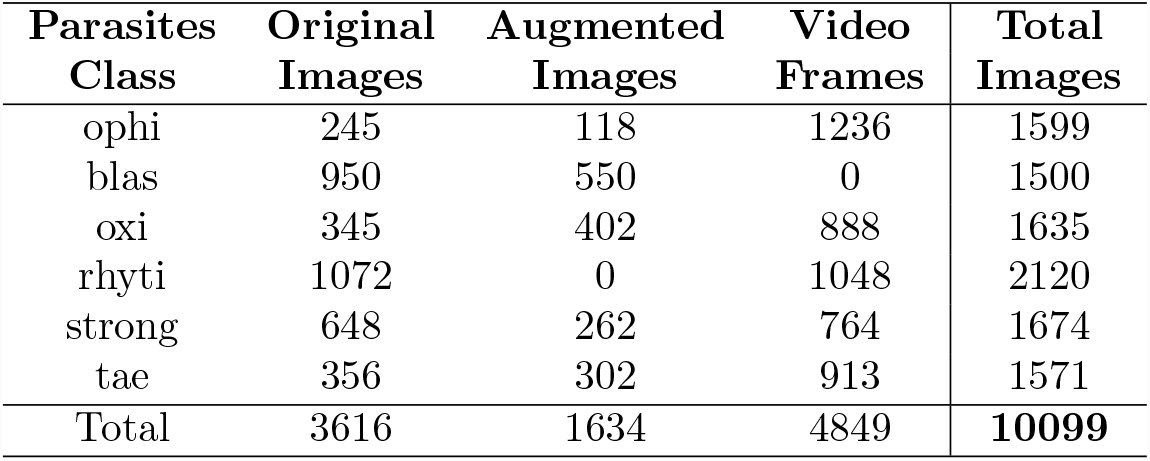
Total number of images in the parasites database after data augmentation

### Region classification using transfer learning with the MobileNet CNN

Transfer learning refers to the ability to reuse an already trained neural network model (referred as pre-trained model) for certain task as the starting point for a model on a second task. In the computer vision domain, pre-trained models are trained on challenging datasets with enormous computing resources.

For instance, MobileNet [23] is a neural network that was trained with the ImageNet database that has about 14 million images. The main characteristic of this CNN is the ability to generate models with performance comparable to other more robust neural networks such as ResNet [34] or VGG16 [35], but with the advantage that it consumes few computational resources thanks to its blocks of depthwise separable convolution blocks (DWSCB) which reduce the calculation time by approximately 8 to 9 times the time of a standard convolution [23]. This makes this model perfect for use on limited resource devices as mobile phones [23], hence its name.

A DWSCB comprises a depthwise convolution (DWC) block and a pointwise convolution (PWC) block with their respective batch normalization layer and ReLU (rectified linear unit) activation function. In summary, a DWC applies a filter to each input channel, as shown in Fig 4, where *M* is the number of inputs and *D*_*k*_ is the kernel size. Then, it feeds these outputs to a batch normalization layer and finally to a ReLU activation function. Subsequently, a 1 × 1 PWC is performed to combine the outputs (see Fig 5), and it again passes through a batch normalization layer and a ReLU activation function. Finally, from Figs 4, 5, and 6, it is possible to notice how a standard convolution operation is factored into a depthwise convolution and a 1 × 1 pointwise convolution operations.

**Fig 4.**
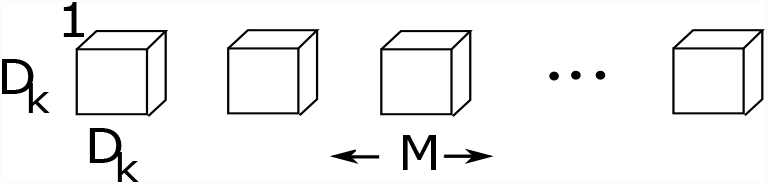
Depthwise Convolution Layer. Adapted from [23].

**Fig 5.**
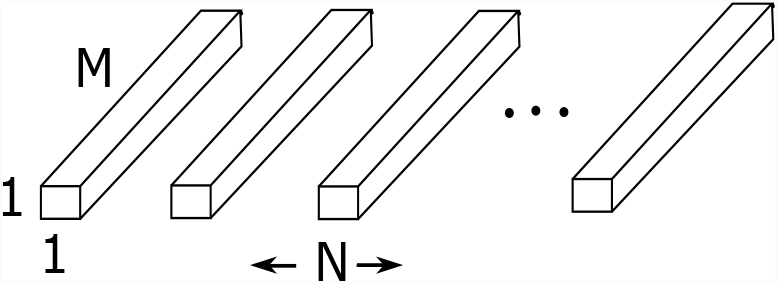
Pointwise Convolution Layer. Adapted from [23].

**Fig 6.**
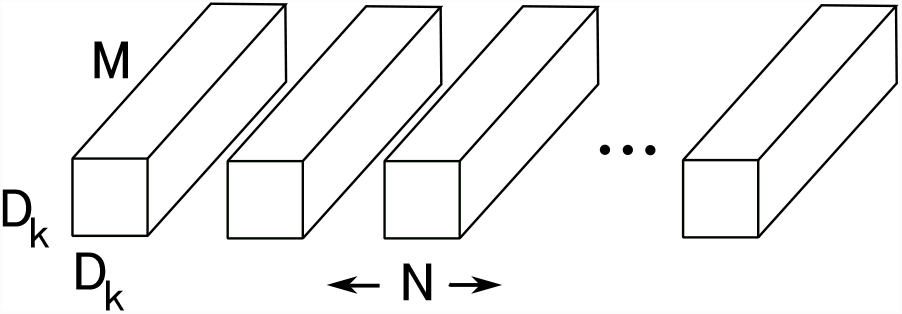
Standard Convolution Process. Adapted from [23].

Afterward, the outputs pass through a standard convolution (SC) block, which filters and combines input parameters into a new set of output parameters. This block again makes use of a batch normalization layer and a layer with ReLU activation function.

Since MobileNet was original designed to classify 1000 classes, we have adapted its architecture to our six classes problem. Hence, we replaced the last classification layer after the average pooling layer with a dense layer of six outputs (one for each parasite class) with a softmax activation function. The adapted MobilNet architecture to our classification problem is shown in Fig 7. From this figure, it should be noted that we included a dropout layer with a dropout rate of 25% to improve the generalization error and to avoid overfitting [24].

**Fig 7.**
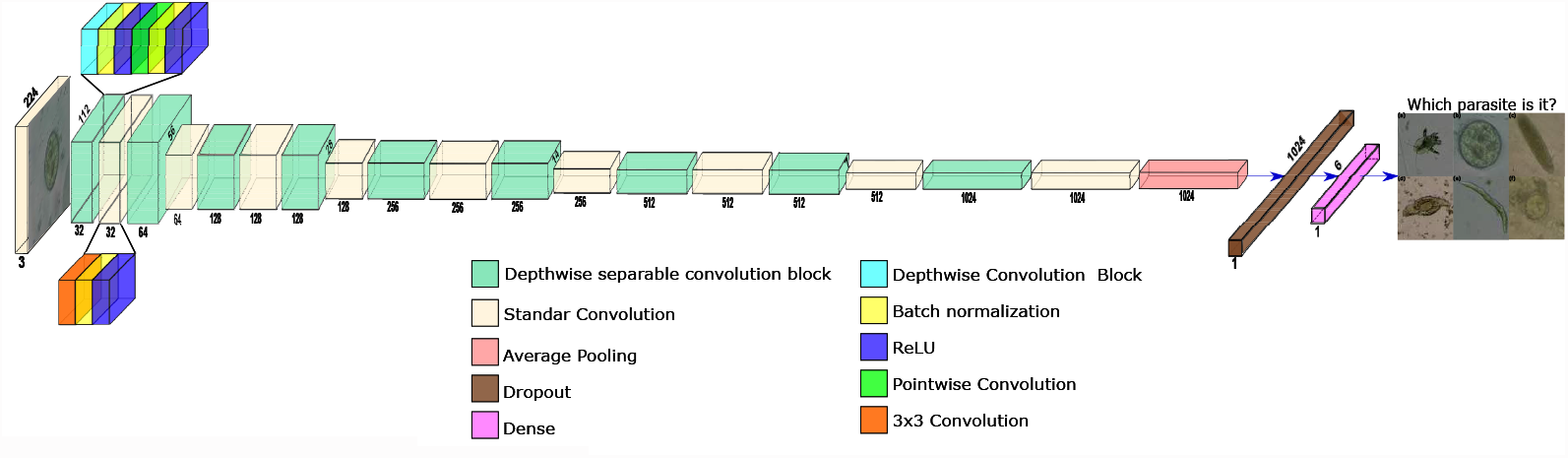
MobileNet Architecture.

### Experimental setup

This section describes the experimentation methodology employed to assess the proposed approach, such as data partition, data augmentation, neural network configuration, comparison with a custom CNN model and the evaluation metrics.

### Data partition

We have used the experimental dataset described in Table 2, which is composed of 10099 instances including original images, augmented images and video frames. We have applied a stratified five-fold cross-validation method [25] (*k* = 5) on the experimental dataset to secure disjoint sets (training and test) and the six parasites classes proportional representation on each fold.

### Neural network configuration

Since we use transfer learning with the MobileNet that was previously trained with different images (i.e., from the Image Net dataset), a parameter tuning process was performed for all layers in the network. Other hyperparameters and settings were stated as:

- Optimization Algorithm: The Adam method was used as it works better than the common stochastic degrading descent. It adapts the learning rate while training for different parameters from first and second-moment estimates of the gradients [27].
- Learning rate: This hyperparameter allows the weights to be updated for each epoch during the training of a neural network. We used a small learning rate as suggested by [26] with an initial value of 0.001. This hyperparameter was reduced by a factor of 0.5 whether the network had not improved its accuracy in 2 epochs.
- Loss function: Since this classification problem is multiclass, the categorical cross-entropy was chosen as the loss function since it leads to faster training as well as improved generalization for classification tasks [36].

### Baseline CNN model

We trained a custom CNN, built from scratch to compare against the transfer learning scheme with MobileNet. This network receives 224×224 images as input and has six outputs classes, similar to the MobileNet network. The custom network has *C* convolutional blocks as shown in Fig 8. Each block is composed of the following layers:

**Fig 8.**
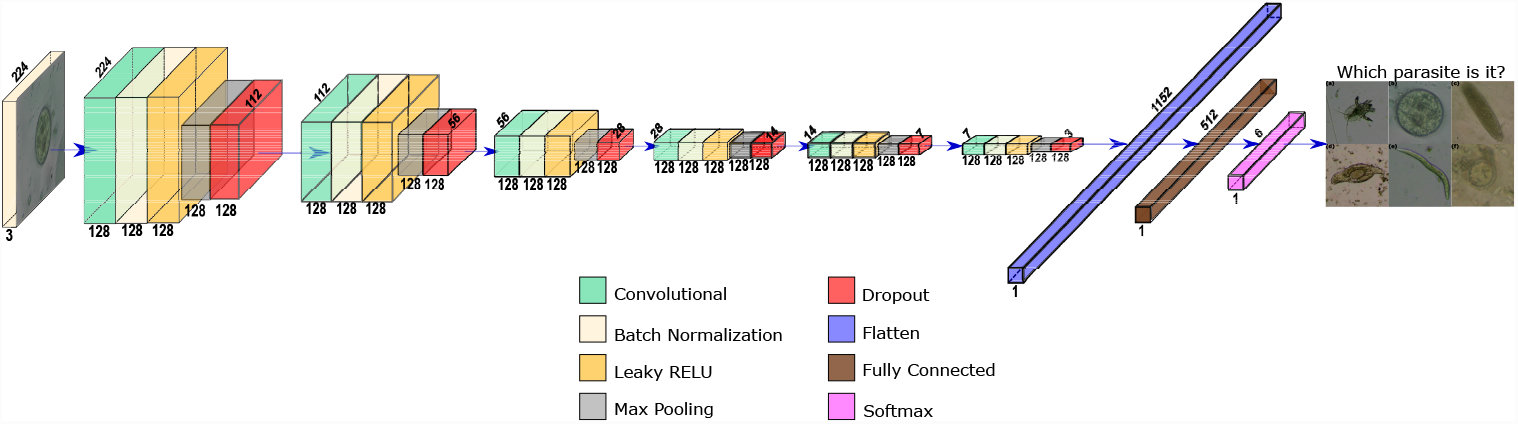
Custom CNN for *C* = 6 blocks and *F* = 128 filters.

- Convolutional layer with *F* filters of size 3 × 3. We set the strides values to 1×1 and zero padding such that the output has the same dimensions as the input.
- Batch Normalization Layer (BN) layer that acts as a regularizer to avoid overfitting and primarily enables training with higher learning rates, which is the cause of faster convergence and better generalization [28].
- Leaky ReLU Layer with a slope of 0.3 has a faster calculation speed and convergence rate, unlike other activation functions such as the sigmoid, thanks to its linearity. Also, this layer was introduced to avoid the vanishing gradient problem since this layer does not cause saturation for negative and positive inputs [29].
- Max Pooling layer allows reducing the dimensionality of the feature maps by summarizing the most active presence of a feature [30].
- Dropout Layer with a drop rate of 0.25 that helps to prevent overfitting [24].

After the convolutional blocks, a flatten layer and one dense layer of 512 nodes with Leaky ReLU activation were added. Finally, the output layer is a six-node dense layer with softmax activation to discriminate each of the six types of parasites.

We varied *C* from 3 to 6 convolutional blocks by keeping constant *F* = 128 filters to explore how depth affects the network performance. Then, we kept constant *C* = 6 convolutional blocks and varied *F* for 32, 64, and 128 filters to analyze the impact of filter size on the network.

### Evaluation metrics

We used the Area Under the Curve (AUC) metric obtained from the ROC (Receiver Operating Characteristic) curve on the same five-fold cross-validation partitions for both the custom CNN and the transfer learning architectures. The ROC curve was obtained by plotting the sensitivity or true positive rate (TPR) on the y-axis against the specificity or false positive rate (FPR) on the x-axis for different threshold decision values varying from 0 to 1.

Moreover, in order to guarantee a fair and statistically reliable comparison, we repeated four times (with different random seeds) the five-fold cross-validation partition scheme, giving a total of 20 runs for each neural network architecture. Since we deal with a multiclass problem, we used the micro-average AUC to compare the transfer learning scheme against the custom CNN. Finally, we calculated the confusion matrix from a five-fold cross-validation run and the overall accuracy of all runs.

## Results and discussion

Before presenting the results with our transfer learning scheme, we are going to present the results of the hyperparameter optimization of the number of convolutional blocks *C* and the number of filters *F* from the custom CNN. In Table 3, we show the average accuracy for different values of *C* and *F* across the 20 runs with different data partitions. We also show the number of trainable parameters for each experiment.

**Table 3.** Average Accuracy and standard deviation (std) with their respective number of trainable parameters and p-values for different values of C and F across 20 runs with different random data partitions. P-values are calculated using 128 filters and 6 convolution blocks as the pivot.

| Filters<br>$F$ | Conv. blocks<br>$C$ | Accuracy<br>(std) | Trainable<br>param | P-value |
| --- | --- | --- | --- | --- |
| 128 | 3 | 95.91 (2.379) | 51,683,334 | 0.06858 |
| 128 | 4 | 95.88 (2.636) | 13,296,006 | 0.12889 |
| 128 | 5 | 94.56 (3.696) | 3,810,054 | 0.07352 |
| 128 | 6 | 95.30 (3.234) | 1,336,454 | — |
| 64 | 6 | 93.68 (4.256) | 485,702 | 1.377e-08 |
| 32 | 6 | 88.69 (5.459) | 198,566 | 0.012 |

Observe that as the number of filters increases, the average accuracy also increases. To select the best custom CNN model, we selected the one with the best average accuracy. In case there are statistically similar results according to the t-test at 95 % accuracy, we chose the one with the lowest number of trainable parameters. According to this strategy, the best model is achieved with 128 filters and 6 convolutional blocks, which hereinafter will be referred as the optimized custom CNN.

The two-dimensional embedding learned by MobileNet before the softmax layer is depicted in Fig. 9. Note that the learned features form quite distinctive clusters, demonstrating that Mobilenet can learn a helpful representation from the images.

**Fig 9.**
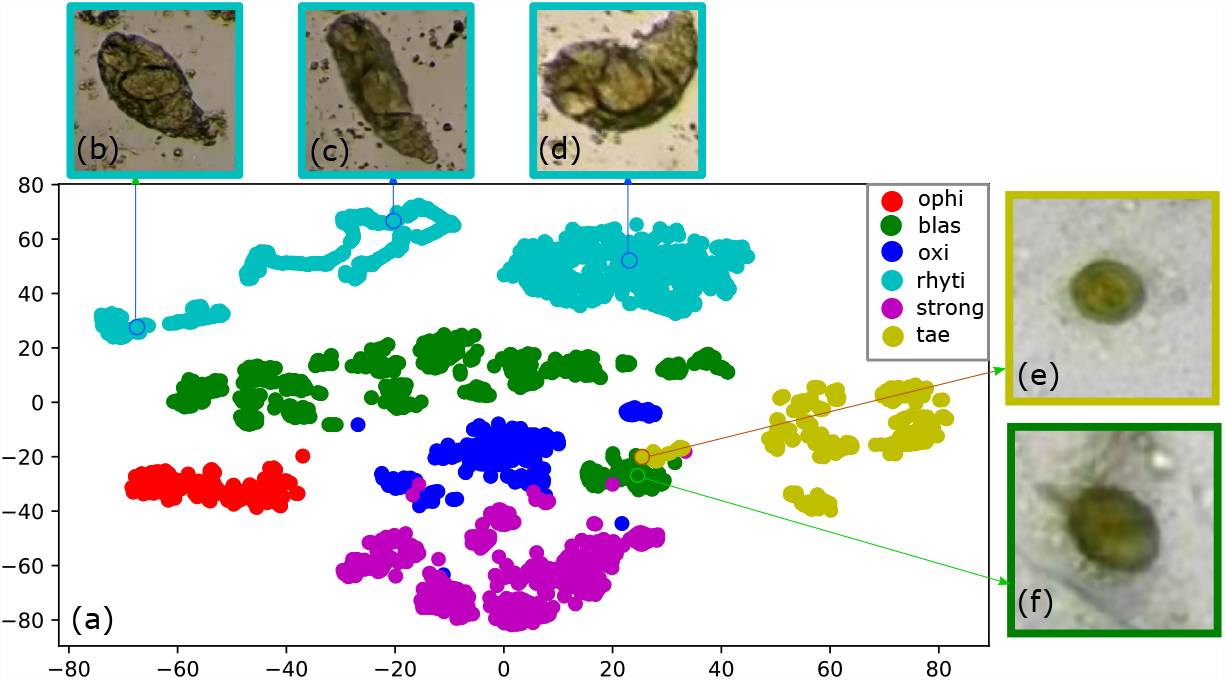
a) The T-SNE embedding shows that Mobilenet is learning a meaningful representation of the image classes. On the other hand, Figs b), c), and d) show representative parasites from three different clusters of the rhyti class. Meanwhile, Figs e) and f) depict some examples of the tae and blas parasites, respectively, from overlapping regions to show their similarity.

It is worth noting that there is some overlapping among classes due to similarities in morphology and color. For instance, observe in the T-SNE embedding how some blas images overlap tae images due to their visual similarities, as depicted in Fig 9e) and Fig 9f) for tae and blas classes, respectively.

Also, note that the T-SNE embedding tends to form three different clusters for the rhyti class. The parasite’s natural motion can explain these three rhyti clusters during the collection stage. To exemplify this, observe the three representative rhyti examples from Fig 9 b), c) and d) taken from these three rhyti clusters. When the parasite was observed by microscope, it might be in a retracted position as shown in Fig 9b. As the parasite’s motion evolves, the parasite might take a more elongated shape as shown in Fig 9c). Finally, this parasite might also be often found in a curved shape which is its feeding position as shown in Fig 9d),. Here, it’s important to note that the neural network could learn this parasite’s motion stages during training.

The T-SNE embedding for the optimized custom CNN can be seen in Fig 10. Although the T-SNE plot tends to group similar parasites, note how the classes overlap far more when compared to MobileNet’s T-SNE. Concretely, classes strong, oxi, and tae are confused with each other more frequently. There is also confusion between tae and blas to a lesser degree.

**Fig 10.**
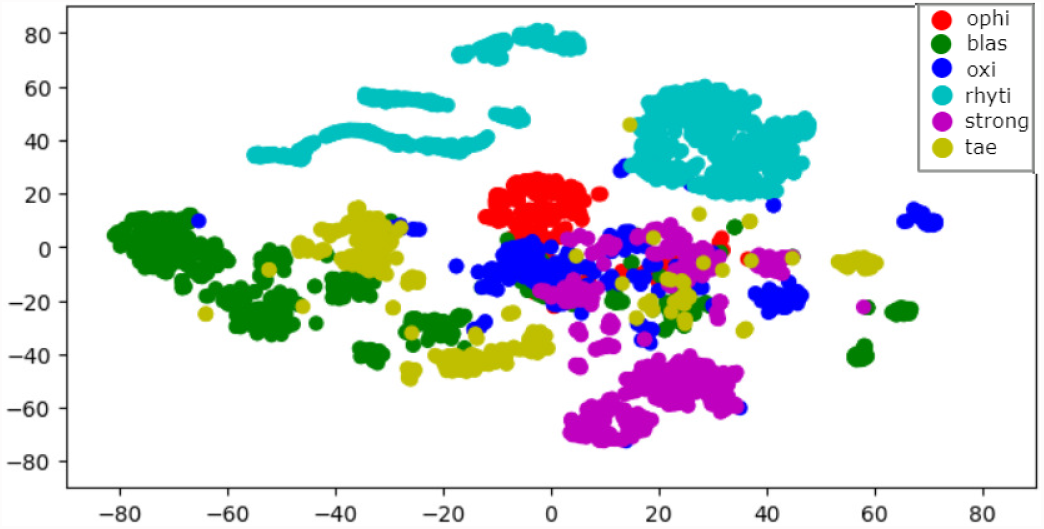
T-SNE plot of the multiclass model for the optimized custom net. Observe how the classes overlap far more when compared to MobileNet’s T-SNE.

The representation of True Positive Rate (TPR) vs. False Positive Rate (FPR) by ROC curves of each class for MobileNet, as depicted in Fig 11, shows that the lowest AUC of 99.919 % is achieved for the oxi parasite and the highest AUC of 100 % for the rhyti parasite. Also, it shows the micro average that adds the contributions of all the classes before calculating the average accuracy, obtaining a value of 99.972 %. The Fig 12 shows similar ROC annotations for the optimized custom CNN, obtaining the lowest AUC of 99.617 % for the oxi parasite and the highest AUC of 99.999 % for the rhyti, and a micro average value of 99.844 %.

**Fig 11.**
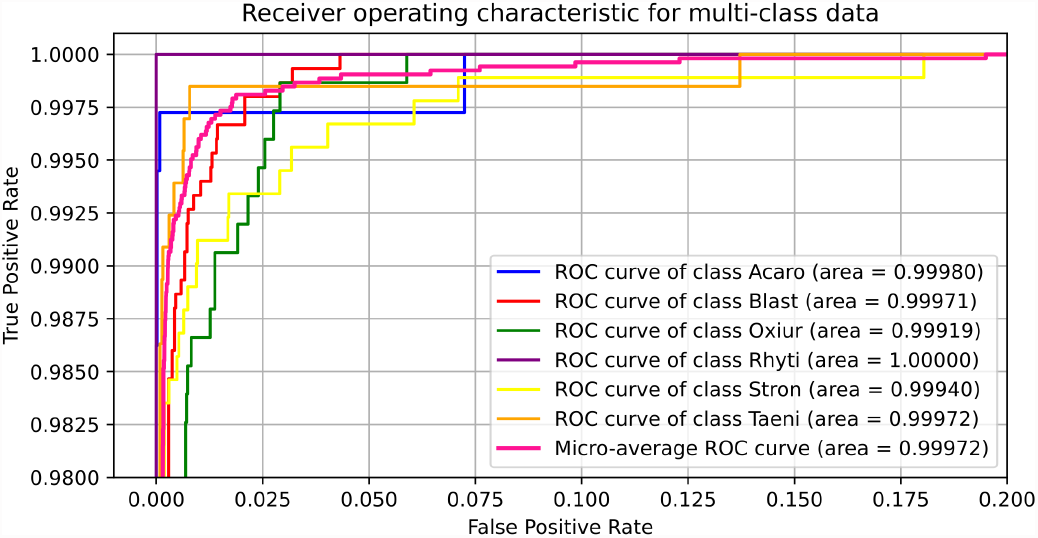
ROC plot for the MobileNet model.

**Fig 12.**
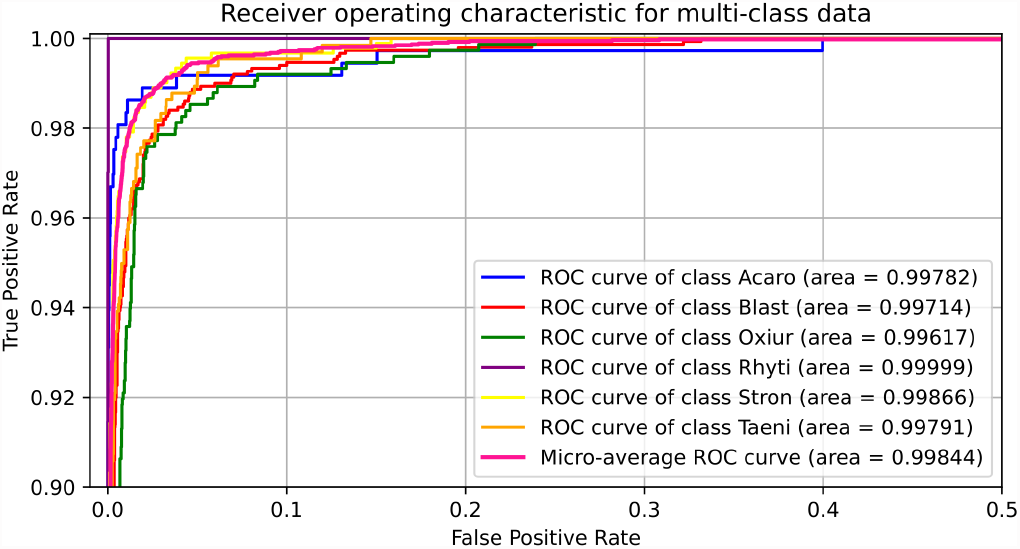
ROC plot for the optimized custom CNN.

In Fig 13 a) we can see the confusion matrix obtained with MobileNet. Looking at Fig 9, it is observed that the classifier effectively distinguishes the classes. Also, a high degree of confusion is observed between the oxi class and the blas and strong classes. Some confusion is also observed between tae and blas classes. This result was expected since both classes overlap in the T-SNE embedding.

**Fig 13.**
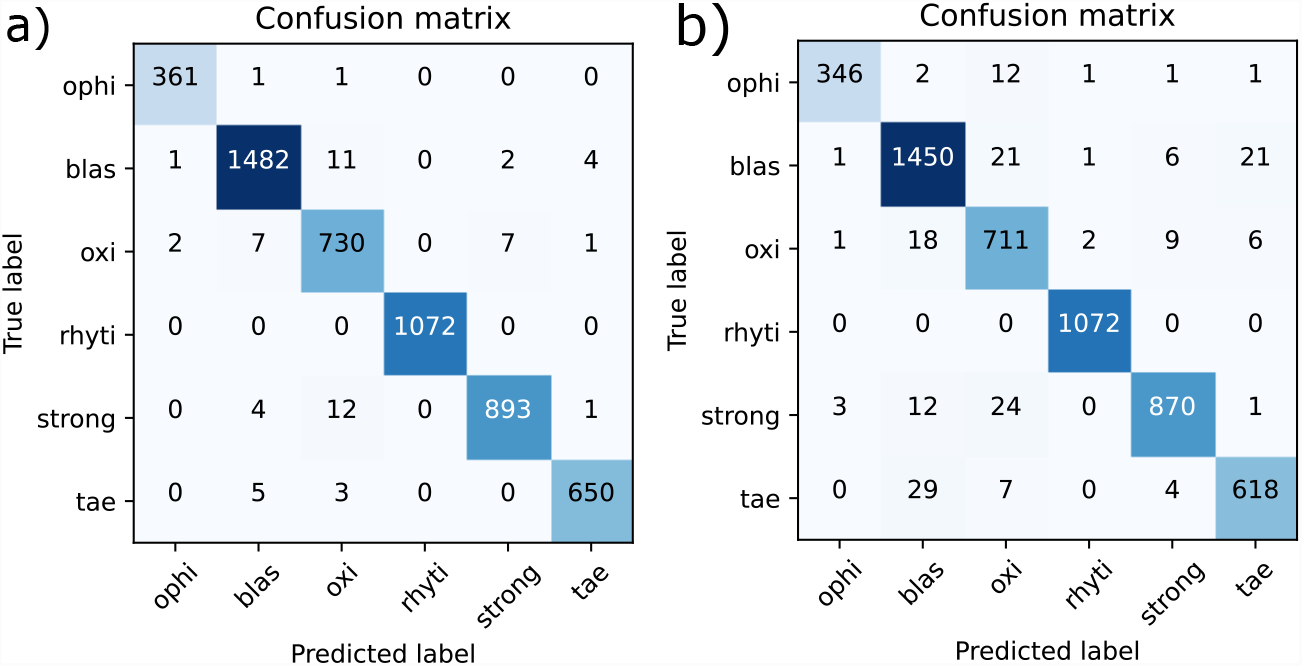
Confusion matrix of the multiclass model by a) MobileNet, b) optimized custom CNN.

Concerning the optimized custom CNN, Fig 13 b) shows its confusion matrix. When compared to the MobileNet, the optimized custom CNN has lower accuracy across all classes. Specifically, the oxi class has a high confusion with the strong and blas classes and moderately with ophi class. Likewise, blas is often confused with strong and tae. In general, there is confusion between all classes except for the rhyti class. This result was expected from what has been seen previously in the T-SNE embedding as shown in Fig 10.

Table 4 shows the accuracy for both the transfer learning scheme with Mobilnet and the optimized custom CNN (i.e., 128 filters and 6 convolutional blocks). Observe that the accuracy value obtained by MobileNet (98.66 %) is better than the optimized custom CNN. We perform a t-test to confirm whether this improvement is statistically significant. The t-test confirms that the mobile network statistically outperforms the optimized custom CNN at a 0.05 significance level, according to the p-value shown in Table 4.

**Table 4.** Accuracy, standard deviation and p-values. The accuracy is the average across the 20 runs with different random data partitions. P-values are calculated using the optimized custom CNN as pivot

| Approach | Accuracy<br>% | Standard<br>Deviation | P-value |
| --- | --- | --- | --- |
| MobileNet | 98.66 | 0.404 | 0.0001304 |
| Optimized custom CNN | 95.30 | 3.234 | - |

Figs 14 a) and b) show the learning curves for the transfer learning scheme and the optimized custom CNN, respectively. With MobiletNet, the neural network achieves better performance before 10 epochs during training, while the optimized custom CNN takes more than 30 epochs. This is because MobileNet, being a pre-trained network, does not learn from scratch as the custom CNN does it.

**Fig 14.**
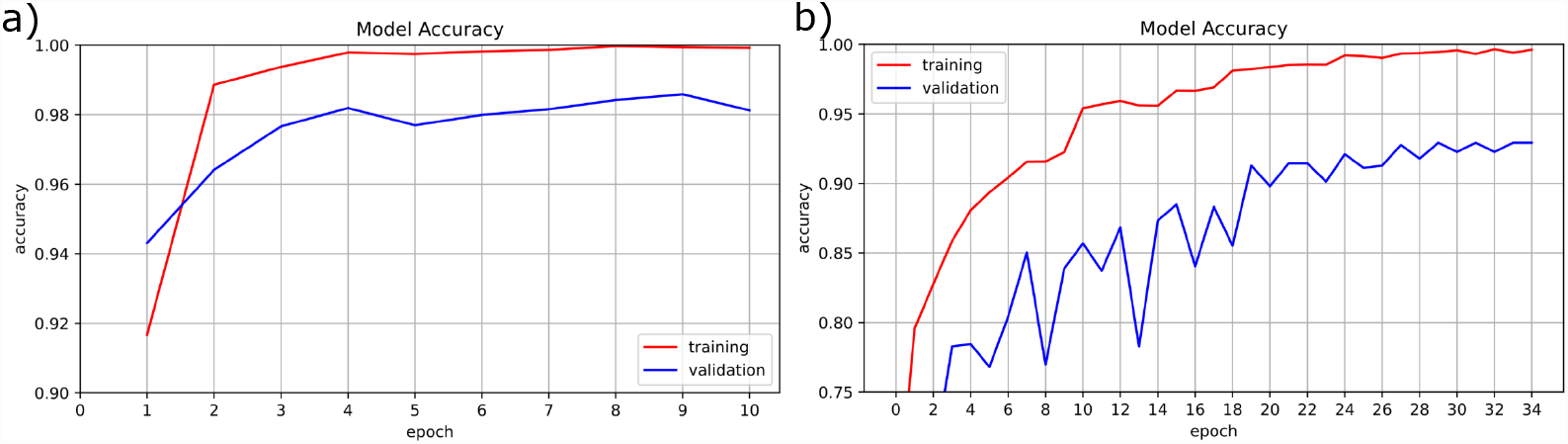
Learning curve for a) MobileNet, b) optimized custom CNN.

### Limitations

Our approach also presents some limitations that decrease its performance due to wrong segmentation or wrong classification predictions. Some wrong segmentation examples are shown in Fig. 15. Concretely, an interior region from a blas parasite was incorrectly segmented in Fig 15a). Moreover, a blas parasite was partially cropped during segmentation in Fig 15b. Finally, the parasite was completely missed in Fig 15c) and instead some debris was captured by the segmentation stage.

**Fig 15.**
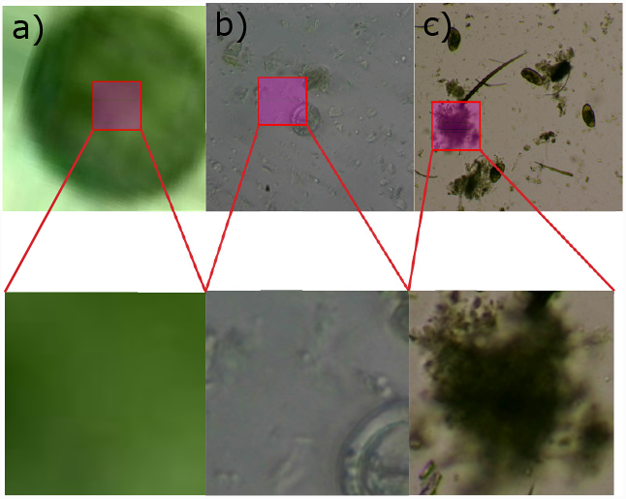
Examples of wrong segmentation

Wrong classification predictions also occurred as shown in Fig. 16, where visually similar parasites are confused. For instance, our approach struggles to differentiate the granules in cytoplasm found on blas parasites, as shown in Fig. 16a), which in turn may cause confusion with the tae class as shown as Fig. 16b). Moreover, our neural network is sometimes not capable of recognizing the tae visual patterns such as its onchosperal membrane as shown in Fig. 16d), which again leads to misclassification as blas. With respect to the oxi parasite from Fig. 16e, it is incorrectly classified as blas since our method is not distinguishing the tae’s oval shape from the blas’ circular shape. In the same figure, it also seems that the similarities between the blas parasite and the tae’s embryon increase the classification error. Finally, the oxy example from Fig. 16g depicts an elongated oval shape which may be confused with the similar shape from the strong larva in movement.

**Fig 16.**
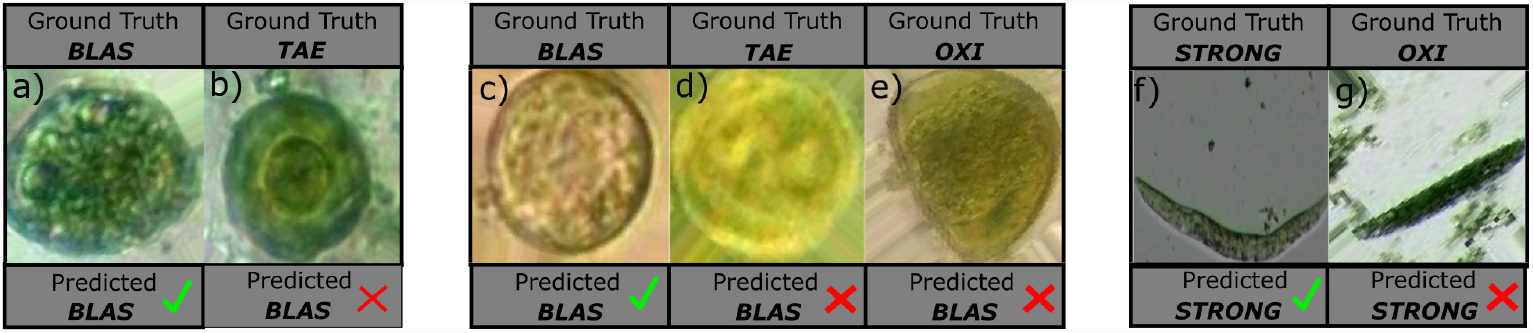
Examples of wrong predictions.

## CONCLUSION

The proposed system for classifying parasitic agents was designed to obtain an adequate performance using a resource-efficient model like MobileNet. Image segmentation mechanisms were a vitally important task since most images had debris or garbage introducing excessive noise due to the nature of their acquisition. The use of data augmentation through video frames for the training stage was crucial to improving network performance due to limited database size. Also, data augmentation algorithms addressed the unbalanced class problem found in this dataset. Our results show that the MobileNet outperforms a optimized custom CNN trained from scratch, demonstrating that transfer learning schemes are suitable to learn relevant features in this domain. Since obtaining labeled data of reptile parasites is an intensive and mainly manual task performed by experts veterinarians, it should be interesting to explore more advanced data augmentation techniques, such as those using the generative adversarial network (GANS) [31]. Moreover, a future study could explore new forms of segmentation (e.g., U-net segmentation [32]) that could improve overall system performance.

## Acknowledgments

The authors thank the Applied Signal Processing and Machine Learning Research Group of Universidad San Francisco de Quito (USFQ) for providing the computing infrastructure (NVidia DGX workstations) to implement and execute part of the developed source code.

## Footnotes

1 https://vivarium.org.ec

## Notes

### Competing Interest Statement

The authors have declared no competing interest.

